# Circadian Clock Control of Muscle Stem Cells Through Temporal Coordination of Notch and Wnt Signaling

**DOI:** 10.64898/2026.06.05.730283

**Authors:** Tali Kiperman, Xuekai Xiong, Jemima Pangemanan, David Horne, Vijay K. Yechoor, Ke Ma

## Abstract

The circadian clock regulates stem cell responses during tissue remodeling and repair. In skeletal muscle regeneration, successful regenerative myogenesis requires a temporally coordinated transition from Notch- to Wnt-driven signaling. However, the mechanisms governing this timing event remain poorly understood. Here, we show that circadian clock activity marks the cycling population of regeneration-activated myogenic progenitors that is induced in concert with Notch signaling. We identify key components of the Notch pathway as direct circadian clock targets and demonstrate that the clock coordinates Notch and Wnt signaling to drive myogenic progression. Genetic activation of the clock in satellite cells, as well as pharmacological clock stimulation, enhanced both proliferative expansion and subsequent differentiation of myogenic progenitors during regeneration. These effects were mediated by early activation of Notch signaling followed by increased Wnt pathway activity at later regenerative stages. Notably, both clock-dependent mechanisms remained functional in dystrophin-deficient mouse muscle and human myoblasts. Furthermore, clock-activating compounds enhanced regenerative myogenesis following acute injury and improved regeneration in dystrophic muscle. Collectively, these findings establish the circadian clock as a temporal regulator of regenerative signaling programs that orchestrate muscle repair with potential for targeted interventions.

## Introduction

The circadian clock exerts critical daily oscillatory control of key biological processes to maintain tissue homeostasis^1–3^. In skeletal muscle, clock function is essential for maintaining tissue mass, its structural integrity and contractile performance^4–9^. The core molecular clock machinery, driven by a transcriptional-translation negative feedback mechanism, has a pervasive effect on skeletal muscle biology via its rhythmic output in gene regulation^10^. Among the rhythmic gene regulations in skeletal muscle, many were involved in metabolic regulation, while genes involved in cell cycle and proliferation and differentiation pathways are also abundant^11^.

Clock-controlled mechanisms in various stem cell compartment are required for tissue homeostasis^2^. In the muscle stem cells (MuSCs), the satellite cells, circadian clock is a key regulatory mechanism that impacts its proliferative and myogenic properties that are crucial for regenerative myogenesis^5,6,12–14^. Many components of the core circadian clock machinery modulate muscle stem cell behavior during regenerative myogenesis, highlighting the concerted clock control of MuSC behavior^5,6,12–14^. The essential core clock transcription activator Bmal1 is required to coordinate the proliferative expansion and myogenic differentiation of muscle stem cells (MuSCs) that facilitate regenerative myogenesis^5,6,15^. In contrast, the clock repressor Rev-erba exerts antagonistic effects in limiting regenerative remodeling^6,16,17^. Furthermore, components of the negative feedback arm of the core clock loop, including Per1/Per2 and Cry1/Cry2, are known to impact myogenesis^13,14^. These studies to date support the role of clock in orchestrating temporal control in myogenic behavior, though the underlying mechanisms involved remain unclear. In addition, despite the current understandings clock modulation of muscle stem cell properties in myogenesis, whether this temporal regulatory control can be targeted to promote tissue remodeling applicable to muscle atrophy or sarcopenic conditions needs to be explored.

The activation, lineage commitment and differentiation of myogenic precursors in skeletal muscle upon muscle injury are essential for regenerative repair to replenish tissue mass^18^. Appropriate clock-controlled muscle stem cell behavior involved in regenerative response thus supports physiological remodeling to counter tissue damage. Previous studies revealed that clock exerts direct transcriptional control of the canonical Wnt signaling pathway, a mechanism underlying clock promotion of myogenic differentiation. Interestingly, a temporally coordinated Notch to Wnt switch is required for proper progression of myogenesis in response to muscle damage, and inappropriate timing of these events leads to deleterious outcomes in regeneration^19,20^. To date, whether the circadian clock, a cell-autonomous timing mechanism, could modulate the Notch to Wnt signaling transition in muscle regeneration is not known.

Muscle stem cell dysfunctions, including impaired asymmetric division during renewal leading to stem cell depletion and compromised myogenic lineage progression, contributes to the regenerative failure in aging and the dystrophic disease^20–22^. Elucidation of mechanisms involved in modulating muscle regenerative capacity could be targeted to restore the deficiency in aged muscle or the dystrophic condition that may mitigate the loss of muscle mass and function with disease progression. Currently, critical gaps still remain to better understand clock function in promoting myogenic repair for muscle disease applications. Studies so far interrogating clock function rely largely on genetic null models, or environmental circadian disruption. Gain-of-function studies are limited, which may provide genetic basis for augmenting clock function for disease applications. The current lack of mechanistic insights into the potential benefits of re-enforcing clock function in disease processes hinders efforts to target the clock for therapeutic interventions.

In the current study, we leveraged a novel genetic clock gain-of-function approach using satellite cell-specific Bmal1 overexpression to explore the function of circadian clock in muscle stem cell regulation. More importantly, development and application of new pharmacological clock activators in acute injury and dystrophic disease conditions provide the mechanistic basis for clock-enhancing interventions for muscle diseases interventions. Applying these genetic and pharmacological tools, the current study elucidated a temporal mechanism linking clock transcriptional control of signaling mechanisms in MuSCs with lineage progression during regeneration amenable to pharmacological targeting to augment regenerative repair.

## Materials and Methods

### Animals

Mice were maintained in the City of Hope vivarium under a constant 12:12 light dark cycle, with lights on at 6:00 AM (ZT0). All experiments were approved by the IACUC committee of the Beckman Research Institute of City of Hope and carried out in concordance with the IACUC approval. Mice with Tamoxifen-inducible Bmal1 overexpression in satellite cells were generated by crossing Pax7-CreERT transgenic mice (*Pax7^tm1(cre/ERT2)Gaka^*/J, Jackson Laboratory Strain: 017763) with a floxed conditional Bmal1 overexpression allele. Conditional ROSA-26 Bmal1 knock-in allele was generated via CRISPR-mediated gene targeting^23^. Mice containing satellite cell-specific conditional Bmal1 overexpression (sBMKI) were further crossed with *mdx* mutant mice (*Dmd^mdx^*, Stock No: 001801, Jackson Laboratory) for three generations to obtain homozygote sBMKI/mdx mice with satellite-cell overexpression of in a dystrophic disease background.

### Cardiotoxin injury and drug delivery

Acute injury was performed by intramuscular injection (IM) of cardiotoxin (CTX, Sigma-Aldrich 217503) into the tibialis anterior (TA), as previously described^6,15^. Briefly, a 60 μL total volume of 10 μM CTX was injected into 3 sites along longitudinal length of TA muscle. 50uL of 2μM Chlorhexidine and CM002, or PBS as control, were injected into TA after injury in C57BL/6 or mdx mice. Injections were performed at 3:00 PM (ZT9) and tissues were collected in the days post injection (dpi) at 1:00 PM (ZT7).

### Primary myoblast isolation and differentiation

Satellite cells were isolated from pooled hind limb muscle of 8-week-old mice of indicated genotypes using MACS isolation^24^ and expanded in culture for primary myoblasts. Briefly, pooled hind limb muscles were collected, finely minced and subjected to collagenase digestion to collect mononuclear cells devoid of multinucleated myofiber. The mononuclear cell digest was washed and seeded on collagen-coated plates, using serial pre-plating to deplete fibroblasts and expand the myoblast population. Myoblasts were subsequently expanded in growth media for 6 passages. Purity of myoblasts obtained were confirmed by uniform differentiation into myosin heavy chain (MyHC)-positive myotubes. Proliferative myoblast cultures were maintained in F-10 medium supplemented with 20% FBS with 2.5 ng/ml bFGF. 2% horse serum (HS)-supplemented DMEM was used for induction of differentiation, as previously described^25^.

### Myoblasts immunofluorescence staining

C2C12 myoblasts (ATCC CRL-1772), Flox control and sBMKI primary myoblasts, or mdx and human DMD primary myoblasts (Creative Bioarray, CSC-C3604) were seeded at 2 × 10^5^/well onto 6-well plates and maintained in growth media until 80% confluency. DMEM with 2% HS was used to induce differentiation together with treatment of the indicated compounds. Myotubes were fixed with 10% neutral-buffered formalin (Sigma, HT501128-4L) on the days indicated after differentiation, permeabilized and blocked with 1% BSA. Primary antibodies were incubated overnight at 4°C, and secondary antibodies applied at room temperature for 1 h. 4′,6′-diamidine-2′-phenylindole dihydrochloride (DAPI, 1 ug/ml) was used to label nuclei. Images were acquired using an Echo Revolve fluorescence microscope. Primary and secondary antibodies used and the dilutions are described in Supplemental Table 1.

### EdU proliferation assay

To quantify the proliferation of myoblast, cells were seeded at 0.5×10^5^/well in 12-well plates and incorporation of 10 μM of 5-ethynyl-2′-deoxyuridine (EdU) was analyzed following addition for 4 to 6 hours before fixation. For *in vivo* analysis, mice were injected intraperitoneally (IP) of 100 μL EdU overnight prior to tissue collection. Detection of EdU was performed using Click-iT EdU Imaging Kit with Alexa Fluor 488 (Invitrogen). DAPI (1 μg/ml) was used for labeling of nuclei. The total number of EdU+ cells was determined using 10 representative fields, and the rate of proliferation was calculated as percentage of EdU^+^/DAPI.

### Skeletal muscle immunofluorescence staining

Muscles were collected and fixed in with neutral-buffered 4% formaldehyde for 24 hours, washed in PBS and incubated in 30% sucrose solution (Fisher Chemical S5-500) for 48 hours prior to embedding in OCT (Sakura Finetek, 25608-930). 10μm cryosections were collected using TA middle portion, fixed with acetone, and permeabilized with TBST containing 0.4% Tween 20. Endogenous IgG was blocked using non-specific mouse IgG (M.O.M Immunodetection kit, VECTOR Laboratories), with further blocking using 5% goat serum (Vector laboratories S-1000) prior to primary antibody incubation at 4^0^C overnight. Secondary antibodies were applied at room temperature for 1 hour and mounted with ProLong Gold antifade reagent with DAPI (Invitrogen P36931). Muscle cross section area (CSA) was calculated from a total of four representative 10X fields from each muscle and plotted for muscle diameter size distribution.

### Hematoxylin and eosin histology

Muscles were collected and fixed with 4% formaldehyde for 72 hours, washed in 70% alcohol prior to embedding. 10μm paraffin-embedded muscle cross sections were processed for deparaffinization, rehydrated, and stained with hematoxylin and eosin.

### Immunoblot analysis

Total protein (20-40 µg) were extracted using immunoprecipitation lysis buffer (3% NaCl, 5% Tris–HCl,10% Glycerol, 0.5% Triton X-10) and resolved on 10% SDS-PAGE gels followed by western blotting on Immuno-Blot PVDF membranes (Bio-Rad). Primary antibodies were incubated overnight at 4^0^C and secondary antibodies applied at room temperature for 1 hour. Blots were developed using Super Signal West Pico PLUS Stable Peroxide Solution (Thermo Scientific 1863095) and Enhancer Solution (Thermo Scientific CN: 1863094), followed by imaging using a chemiluminescence GE BioSciences AB Amersham Imager 680. Primary and secondary antibodies used and the dilutions are described in Supplemental Table 2.

### RNA extraction and RT-qPCR analysis

Trizol (Invitrogen) or RNeasy miniprep kits (Qiagen) were used to isolate total RNA from snap-frozen tissues or cells. Reverse transcription was carried out using a Revert-Aid RT Reverse Transcription Kit (Thermo Fisher K1691) and ran on a SimpliAmp Thermal Cycler (Thermo Fisher A24811). Quantitative PCR was performed using SYBR Green Master Mix (Thermo Fisher A25742) and ViiA 7 Real-Time PCR System (Applied Biosystems). Relative expression levels of target genes were determined using the comparative Ct method and normalized to 36B4 as internal control. Primer sequences are listed in Supplemental Table 3.

### RNA-seq analysis

RNA-seq reads were aligned with STAR software to the mouse reference genome mm10, and unique reads were quantified using HTSeq-count with GENCODE annotations. The RNA-seq reads were normalized and log-transformed using limma edge R packages. *P*-values and the false discovery rate (FDR) were calculated from raw counts using DESeq2. Fold-change > 1.5, 50% fragments per kilobase of transcript per million mapped reads (FPKM) >0.1, unadjusted *P*-value <0.05, and FDR < 0.25 were used as cut-off for differentially expressed genes. Global analysis heatmaps were produced using heatmap.3 and the gplots package in R, using log2 (FPKM + 0.1) values. The Pearson dissimilarity (1–Pearson correlation coefficient) was used as the distance metric for hierarchical clustering of rows and columns. Expression was centered so that each gene had a mean of 0, and centered expression was capped at –3 and 3, for clustering and visualization.

#### Analysis of single-cell RNA sequencing datasets

Published scRNA-seq datasets GSE143437, GSE159500 and GSE162172 and GSE232106 containing analysis of dpi 0 to 7 of muscle regeneration were used in the analysis^26,27^. Marker genes for each author-defined myogenic cell state were identified with Scanpy rank_genes_groups (Wilcoxon rank-sum, BH-adjusted). A focused differential-expression comparison of non-cycling versus cycling MPCs was performed (Wilcoxon). Arntl (Bmal1), CLOCK, Notch module and canonical Wnt module values were summarized as violin plots across the ordered cell states, and directly compared between non-cycling and cycling MPCs by two-sided Mann-Whitney U test.

### Chromatin Immunoprecipitation (ChIP) analysis

C2C12 cells were collected, crosslinked and sonicated to shear the chromatin according to manufacturer’s instructions of the Simple ChIP Plus Sonication Kit (Cell Signaling Technology 56383). Immunoprecipitation was performed with 5μg Bmal1 antibody (AB93806; Abcam) or rabbit IgG. The immunoprecipitated chromatin fragments were processed and purified using the Simple ChIP Plus Sonication Kit and DNA Purification Buffers (Cell Signaling Technology 14209S). Real-time PCR using Taq Pro Universal SYBR qPCR Master Mix (Vazyme Q712-03) was carried out with specific primers flanking the identified E-box sites using genomic primers for Tbp promoter as negative control. Primer sequences are listed in Suppl. Table 4. Data are expressed as fold enrichment over IgG after normalization to 1% input.

### Statistical analysis

Experiments results were analyzed and graphed using PRISM, and data were presented as mean + SD. Each experiment was repeated at least twice. Biological replicates were indicated for each experiment in figure legends. Unpaired two-tailed Student’s t test was used for tow group comparisons, and one-way ANOVA for multiple comparisons with Tukey’s post hoc analysis between two groups were performed as appropriate. P<0.05 was considered statistically significant. GraphPad Prism software (version 6.01) was used for generating graphs, and BioRender was used for schematic illustrations.

## Results

### Circadian clock marks cycling myogenic precursor cells in muscle regeneration and exerts direct transcriptional control of the Notch signaling pathway

Circadian clock is known to drive stem cell response to tissue remodeling^1,2^. In skeletal muscle, the regenerative response induced by acute injury is an established paradigm to study muscle stem cell functions. To explore potential dynamic regulation of clock activity during muscle regeneration, we surveyed core clock genes using previously published sc-RNA-seq datasets that delineated myogenic progression in response to muscle damage at days post injury (dpi) 1 to 7^26,27^. Their expression in all cell types involved in regeneration (Suppl. Fig. S1A) were plotted, along with distribution in subgroups of MuSCs and myogenic progenitor cells (MPCs, Suppl. Fig. S1B). Interestingly, though neither Bmal1 or CLOCK was robustly expressed in mature myonuclei, they were enriched in MuSCs (Fig. S1B). Furthermore, these transcriptional activators that drive clock oscillation displayed strong enrichment among subpopulations of cycling and differentiating vs non-cycling MPCs (**Fig. 1A**). By applying a Pseudotime analysis to trace the lineage trajectory of MuSC to committed and differentiated MPCs, we found that Bmal1 and CLOCK expression displayed a regulatory dynamic along regenerative myogenesis that were maintained at markedly elevated levels in MuCSs with a sharp return to basal suppressed states during lineage commitment and differentiation (**Fig. 1B**). It is intriguing that this dynamic regulation aligned closely with Notch gene expression score, a key signaling pathway in determining MuSC proliferative expansion and renewal. We therefore determined the distribution of Bmal1 and CLOCK together with key signals driving MuSC behavior in regeneration (Suppl. Fig. S2). Their aggregate expression among MuSCs and MPCs steadily increase since day 2 of regeneration (Suppl. Fig. S2A). Among distinct myogenic subclusters (Fig. S2B & S2C), the levels of both core clock regulators were induced in MuSC and cycling MPCs, with gradual dampening of expression along commitment and differentiation of MPCs (**Fig. 1C & S2D**). Notably, this regulation mirrored that of the composite Notch pathway score, and was partially correlated with Wnt score among MuSC and MPCs, though Wnt was maintained at high levels in mature myonuclei. This selective and intricately controlled induction of clock in MPCs actively participating in regeneration closely correlated with Notch signaling activation, suggesting a specific function in this process.

**Figure 1.**
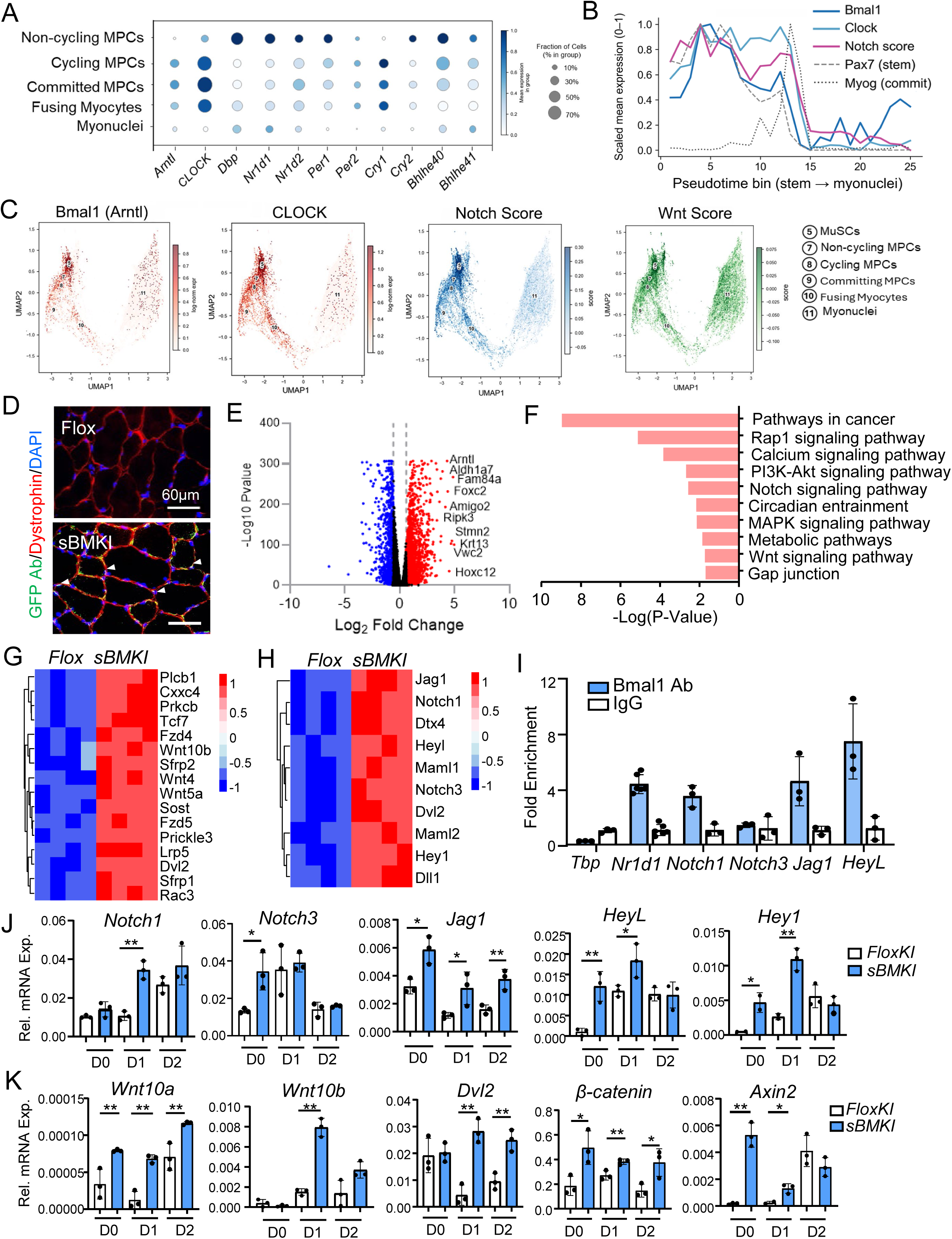
Circadian clock direct transcriptional control and functional regulation of the Notch and signaling pathway. (A-C) Identification of molecular clock activation in muscle stem cell populations from sc-Seq analysis during muscle regeneration. Published scRNA-seq datasets GSE143437, GSE159500 and GSE162172 and GSE232106 were used in the analysis. (B) Pseudotime trajectory analysis across the ordered myogenic states (MuSCs 1–6 → non-cycling/cycling/committing MPCs → fusing myocytes → myonuclei), with clock gene expression together with stem cell (Pax7) and committed MPCs markers (Myogenein) shown. Notch score was included for the analysis. (C) Bmal1 and CLOCK enrichment within MPCs subgroups along lineage progression in regeneration with correlation of Notch and canonical Wnt signaling scores. (D) Representative immunofluorescence staining using GFP antibody (green) with Dystrophin (red) of Flox control and sBMKI TA muscle. Scale bar: 60 μm. Arrowheads denote satellite cells with GFP expression. (E, F) Volcano plot representation of differentially expressed genes (E), and KEGG pathway enrichment analysis (F) of isolated sBMKI primary myoblasts as compared to Flox controls (n=3/group). (G, H) Heatmap of significantly expressed genes in Flox and sBMKI primary myoblast involved in NotchE), or the Wnt (F) signaling pathways. (I) ChIP-qPCR analysis of chromatin occupancy of Bmal1 on identified E-box sites of proximal promoter regions (+2Kb from TSS) of genes involved in Notch signaling. E-box site on Nr1d1 gene was used as a positive control. N=3-6/group. (J, K) RT-qPCR analysis of dynamic expression of components of Notch (H), and Wnt (I) pathways in in Flox and sBMKI primary myoblast before and following myogenic differentiation for 2 days. N=3/group. * and **: p<0.05 or 0.01 by Student’s t test.

To experimentally determine clock activation of stem cell functions in skeletal muscle, we established a Bmal1 conditional overexpression model with tamoxifen-inducible clock gain-of-function in satellite cells (sBMKI), by crossing a Floxed Bmal1 allele (FloxKI), containing flox-stop-flox sequence flanking a Flag-tagged cDNA with bicistronic GFP, with the Pax7-CreERT2^28^ mice^23^. 5 days following tamoxifen induction, the Flag-tagged Bmal1 expression was detected in skeletal muscle groups examined together with GFP (**Fig. S3A**), but not in the liver or heart. Notably, Bmal1 overexpression as detected by a GFP antibody was found in satellite cells residing outside sarcolemma with dystrophin co-immunostaining (**Fig. 1D**). We first performed transcriptomics profiling of satellite cell-derived primary myoblasts from sBMKI and Flox controls to interrogate clock-controlled outputs in modulating satellite cell properties. Global analysis of differentially expressed genes revealed enrichment of up-regulation of 2659 genes comprising 16.6% of all detected transcripts, as compared to 1022 (6.4%) displaying down-regulation (**Fig. 1E**), as expected of Bmal1 gain-of-function in these cells. Bmal1 (Arntl) was among the top up-regulated genes with a 4.4-fold induction. Pathways in cancer, circadian entrainment and metabolic regulation^11^, known to be clock-controlled outputs, were identified among the top up-regulated biological processes in sBMKI myoblasts revealed by KEGG pathway analysis (**Fig. 1F)**. Notch and Wnt signaling, major mechanisms that drive myogenic precursor renewal and myogenic lineage progression during muscle regeneration^19,29^. Both pathways were found to be enriched by the pathway analysis, and their key signaling components were markedly induced (**Fig. 1G & 1H)**. This finding implicated the functional control of Wnt and Notch pathways by clock in MuSCs that may impact their activation and myogenic properties. The Wnt pathway is under the clock transcriptional control in myogenic progenitors, as specific components were identified as direct Bmal1 targets^5,30^. To dissect whether genes involved in Notch signaling could be transcriptional targets of clock, we surveyed E-box elements that meditate Bmal1 chromatin binding within the proximal promoter regions of candidate targets^31^. Using the known Bmal1 target *Nr1d1* promoter as a positive control, ChIP-qPCR analysis revealed that among these putative targets, Notch1, Jagged 1 (Jag1) and HeyL promoter E-boxes displayed association with Bmal1 (**Fig. 1G**), while Notch 3 showed minimal binding affinity. Furthermore, these Notch signaling genes, Notch 1, Jag1 and HeyL, were robustly induced in sBMKI myoblasts as compared with Flox controls during normal proliferative growth or myogenic differentiation (**Fig. 1J**). Importantly, most Notch signaling components examined in sBMKI cells were modulated similarly as controls following day 2 of myogenic differentiation, suggesting a normal dynamic regulation to myogenic stimuli. Similar up-regulations of Wnt pathways genes that were identified as Bmal1 targets^5^, including Wnt10a, Wnt10b, Dvl2 and β-catenin, were found in sBMKI myoblasts under these conditions (**Fig. 1K**). Notably, Axin2 expression level, indicative of the transcriptional activity of the Wnt cascade, was robustly up-regulated early before and at first day of differentiation, though was effectively repressed to a comparable level as that of the controls at day 2.

Under normal physiological conditions without injury, the overexpression of Bmal1 in satellite cells did not elicit significant impact on normal muscle growth or function. Tissue weight of distinct muscle groups, Tibialis Anterior (TA) and gastrocnemius (GN), were comparable with Flox controls (**Fig. S3B & S3C**). H/E histology (**Fig. S3D**), and laminin staining to assess myofiber cross section area (**Fig. S1E**) revealed comparable fiber size. Muscle function measured by grip strength were also similar between these groups (**Fig. S3F**).

### Genetic clock gain-of-function in satellite cells prolonged the myogenic phase with a delayed peak in muscle regeneration

Muscle regeneration follows a time course driven by satellite cell proliferative expansion after activation that progresses to nascent myofiber development and fusion^18^. To determine the functional properties of MuSCs with ectopic expression of Bmal1, we subjected the sBMKI mice to acute injury by cardiotoxin and examined the dynamics of regenerative response in flox control and sBMKI mice for 30 days post injury (dpi 30). Acute muscle injury in the TA muscle elicited a strong response of circadian clock and regenerative myogenesis by MuSC clock gain-of-function. As expected, Bmal1 protein was elevated at basal state without injury in sBMKI TA muscle. Both Bmal1 and CLOCK protein steadily rose after injury along the time course of regeneration, with marked inductions at dpi5 to 30 (**Fig. 2A**). In contrast to the endogenous Bmal1 protein that was induced during regeneration peaking at dpi 3-5 followed by a return to basal levels at day 30, the high levels of Bmal1 and CLOCK protein were likely due to persistent myofiber overexpression of Bmal1 following regeneration. Further analysis revealed an interesting regulatory dynamic of the myogenic response observed in the sBMKI mice (**Fig. 2B**). At the early regenerative stage of 3 days after injury, the myogenic response, as indicated by the induction of Pax7, Myf5, MyoD and Myogenin gene expression, displayed a modest attenuated tendency in sBMKI mice as compared to controls. In contrast, at later time points of regeneration from dpi5 to 14, their expression levels were mostly elevated. 30 days after injury when regeneration was completed, the expression of these genes in sBMKI mice returned to comparable levels as in the Flox controls. Examination of the protein levels of these myogenic regulators further validated the gene expression analysis (**Fig. 2C**). The reduction of Pax7, Myf5 and Myogenin protein were evident at dpi3 and 5 in sBMKI mice. However, at later stages of regeneration of dpi 14 as well as dpi30, when the early induction of these factors returned to undetectable levels in the controls, as expected of normal regeneration, these proteins remained elevated in sBMKI mice. These findings revealed a distinct regenerative trajectory in sBMKI mice, with depressed level of myogenic induction at early stage of regeneration and a persistent myogenic drive beyond the normal response, suggesting a delayed, yet prolonged regenerative phase in mice with MuSC clock overexpression. We further determined the progression of myofiber size via laminin immunostaining to examine nascent myofiber development during specific times of regeneration. Following CTX injury at dpi5 in sBMKI mice, there was an increase in the number of myofibers that display smaller diameter as compared to the controls, as indicated by the quantitative analysis of their cross section area distribution (**Fig. 2D**). A marked shift of myofiber size toward 500µm^2^ or smaller was indicative of nascent regenerated fibers at this early stage of regeneration. In comparison, by day 10 and 14 of regeneration, there was a significant shift of myofiber size in sBMKI mice from the smaller populations with a tendency toward larger size range (**Fig. 2E**). By dpi 30 when the regeneration is largely completed in normal regeneration, sBMKI TA displayed reduction of myofibers at the smaller end of the size distribution (500-1,000 µm2), while the abundance of larger fibers of ≥2,000 µm2 diameters were significantly elevated (**Fig. 2F**). These analyses together indicate that clock gain-of-function in MuSCs resulted in a tempered early regenerative response, though regeneration persisted with completion at a delayed time point.

**Figure 2.**
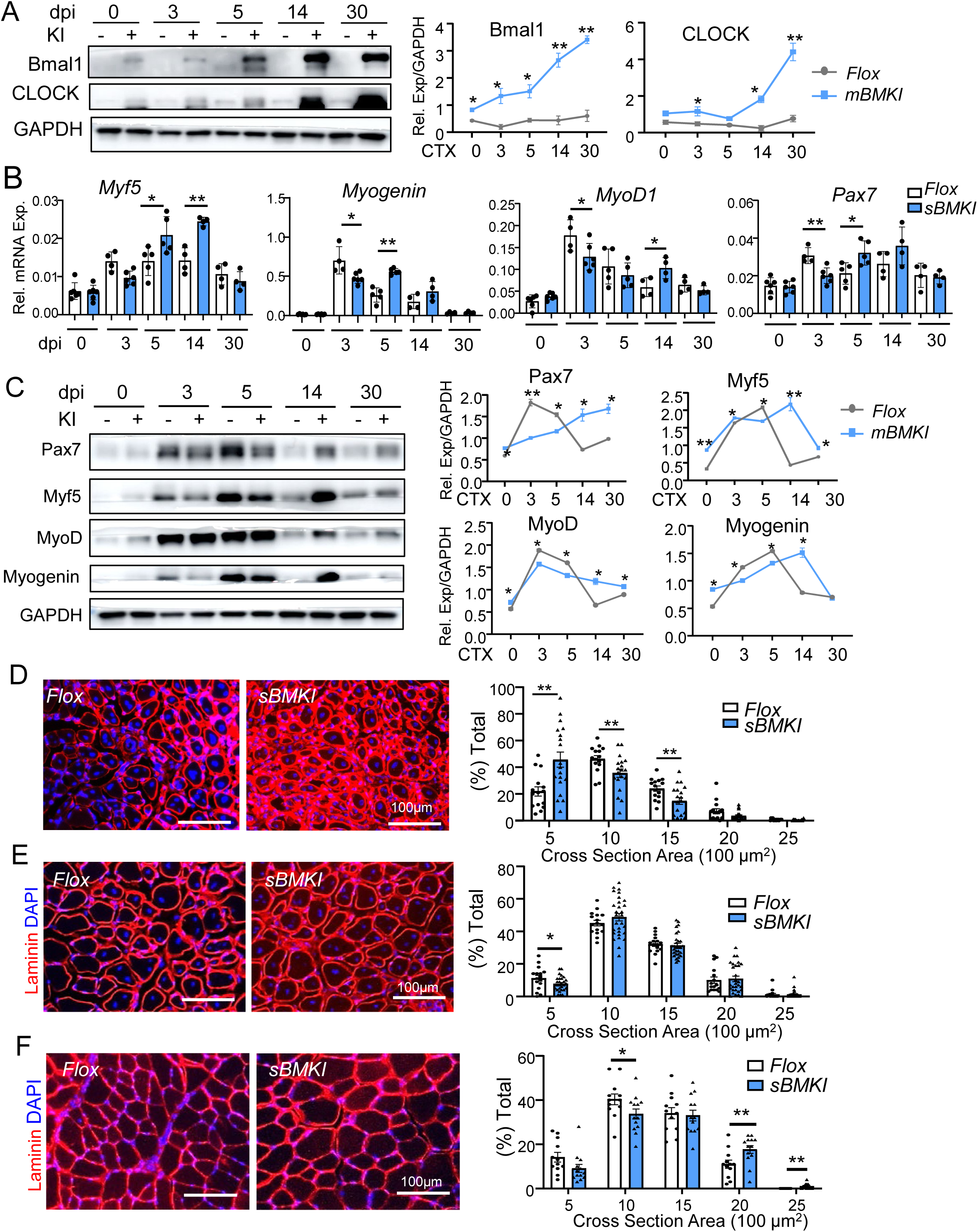
Satellite cell clock gain-of-function prolongs the regenerative myogenic phase of muscle regeneration. (A- C) Gene expression analysis of the dynamic myogenic response along the regeneration time course following cardiotoxin injury of TA muscle. Representative immunoblot analysis of clock protein induction (A), myogenic response (C), and RT-qPCR analysis (B) of Flox and sBMKI mice at indicated times till 30 days after injury. dpi: days post injury. For immunoblot analysis, each lane represents a pooled sample of 5-7 mice/group, with quantification of three repeats. (D-F) Representative images of laminin immunofluorescence staining of myofiber cross sections at dpi 5 (D), dpi 14 (E) and dpi 30 (F) with corresponding quantitative analysis of myofiber section area. Scale bar: 100μm.

### Clock gain-of-function in MuSCs promotes satellite cell proliferative expansion and myogenic progenitor properties

We next examined the early regeneration phase in sBMKI mice via determination of the nascent myofiber development using eMyHC immunofluorescence staining to identify new myofiber formation. Despite the attenuated early myogenic response in sBMKI mice (**Fig. 2 & Fig. S4**), examination of eMyHC^+^ levels at dpi 5, past the normal peak of new myofiber generation, revealed the abundance of new eMyHC^+^ myofibers in sBMKI mice, suggesting a delayed peak of nascent myofiber formation in line with the delayed myogenic response (**Fig. 3A**). Furthermore, through EdU labeling with Pax7 co-staining, satellite cell proliferation at day 7 dpi near the end of normal expansion revealed increased number of proliferative satellite cells in sBMKI mice, indicative of a prolonged satellite cell expansion phase (**Fig. 3B**). A temporal progression from Notch pathway activation at early muscle regeneration to Wnt signaling at later stages is required for normal regenerative myogenesis^19^. Based on our prior findings of clock transcriptional regulation of signaling components of the Wnt cascade^5,30^ together with current results of its regulation of the Notch pathway, we postulated that clock coordination of the temporally controlled Notch and Wnt signaling may underly the delayed yet prolonged regenerative response in sBMKI mice. Analysis of both pathways along the regeneration time course revealed that a major Notch receptor in MuSCs, *Notch1*, and Notch signaling downstream effector *HeyL* were induced at early stages of regeneration, through they were effectively suppressed at the completion of regeneration at day 30 similarly as the controls (**Fig. 3C**). In contrast, a known Bmal1 targets within the Wnt pathway, Wnt10a, was induced at day 5 and 14 of regeneration, while expression of the Wnt receptor Fzd5 was attenuated at 3 days following injury but was robustly elevated at dpi 14 that persisted at day 30 (**Fig. 3D**). Thus, forced clock expression in MuSCs modulates the temporal progression of these signaling events driving myogenesis with enhanced Notch activity early in regeneration together with a prolonged Wnt pathway activation. Given the critical function of Notch activation in MuSC self-renewal and proliferation in muscle regeneration, we explored whether these changes induced by Bmal1 overexpression underlies the proliferative phase of regenerative repair and isolated satellite cells at 5 days after CTX injury to determine their activation and myogenic commitment properties (**Fig. 3E & 3F**). Identification of proliferative satellite cells via EdU/Pax7 labelling revealed a ∼27% increase of proliferative rate among these cells isolated from sBMKI mice as compared to that of controls (**Fig. 3E**). Furthermore, a significantly higher proportion of cells double positive for MyoD and Myf5 indicative of activated myogenic progenitors^32^ were found in the satellite cells from sBMKI mice (**Fig. 3F**).

**Figure 3.**
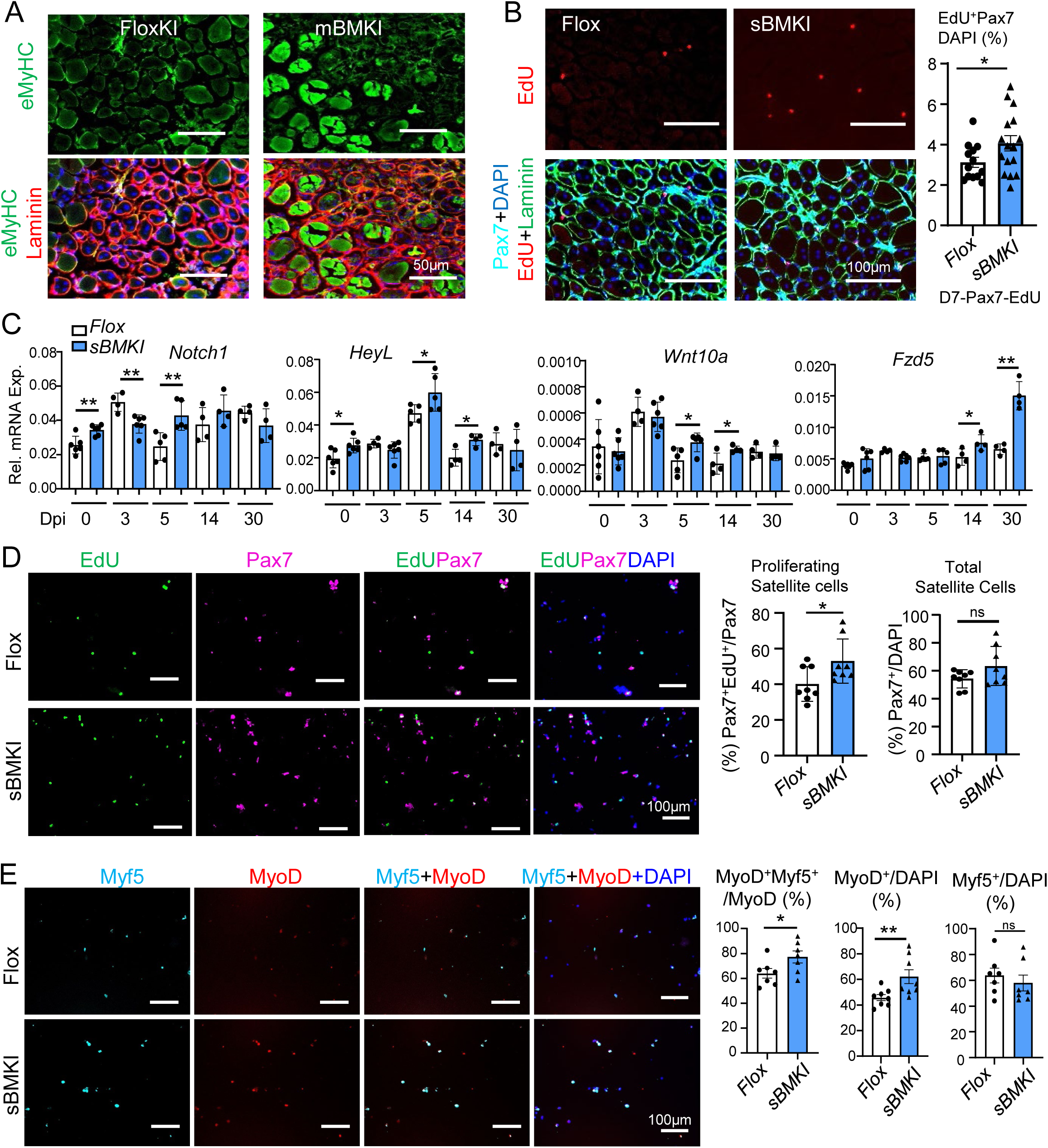
Clock promotes satellite cell proliferative expansion and myogenic progenitor function. (A) Representative images of immunofluorescence staining of myosin heavy chain (MyHC, green) and laminin (red) to assess nascent myofiber formation in Flox and mBMKI mice at dpi5. (B) Representative images of Pax7 immunofluorescence staining (green) with EdU fluorescence (red) and the quantitative analysis of proliferating satellite cells in TA muscle of Flox and sBMKI mice 7 days after CTX injury. (C) RT-qPCR analysis (B) of Notch and Wnt pathway genes in Flox control and sBMKI mice at indicated times along the regeneration time course. N=5-6 mice/group. * and **: p<0.05 or 0.01 by Student’s t test. (D & E) Representative images of immunofluorescence staining of Pax7 (ed) with EdU (Green) fluorescence labeling to assess proliferation of satellite cells (D), and MyoD (red) or Myf5 (Cyan) to assess MuSC properties (E) using satellite cells isolated from Flox and sBMKI mice at dpi 5 after CTX injury (n=5/group). Scale bar: 100μm.

We also amplified satellite cells via ex vivo expansion resulting in activated primary myoblasts. Under normal proliferative growth condition, these satellite cell-derived sBMKI myoblasts displayed a significant ∼38% increase in proliferative rate than that of the controls (**Fig. 4A**). Notably, while as the rate of proliferation of Flox controls dropped following induction of myogenic differentiation using 2% horse serum, as expected, the elevated level of proliferation of sBMKI cells was maintained, a finding that was in line with the observation of delayed but prolonged myogenic response in vivo. Furthermore, analysis of cell cycle regulators revealed induction of Cyclin D1 and Cdk4 genes in proliferative sBMKI myoblasts with their expression effected inhibited by day 2 of differentiation (**Fig. 4B**). Most interestingly, despite the up-regulated cell cycle regulators, induction of Myod1, Myogenin and eMyHC genes were elevated in these cells, suggesting enhanced myogenic differentiation (**Fig. 4C**). To test whether increased proliferation of sBMKI cells was mediated by Notch signaling, they were treated with a specific inhibitor CB103 that blocks the transcriptional activation of Notch pathway. A relatively low concentration of 2uM of CB103 was sufficient to abolish increased rate of proliferation of sBMKI cells down to a comparable level observed in Flox controls (**Fig. 4E**), suggesting that Notch activity indeed mediated the proliferative phenotype induced by Bmal1 overexpression.

**Figure 4.**
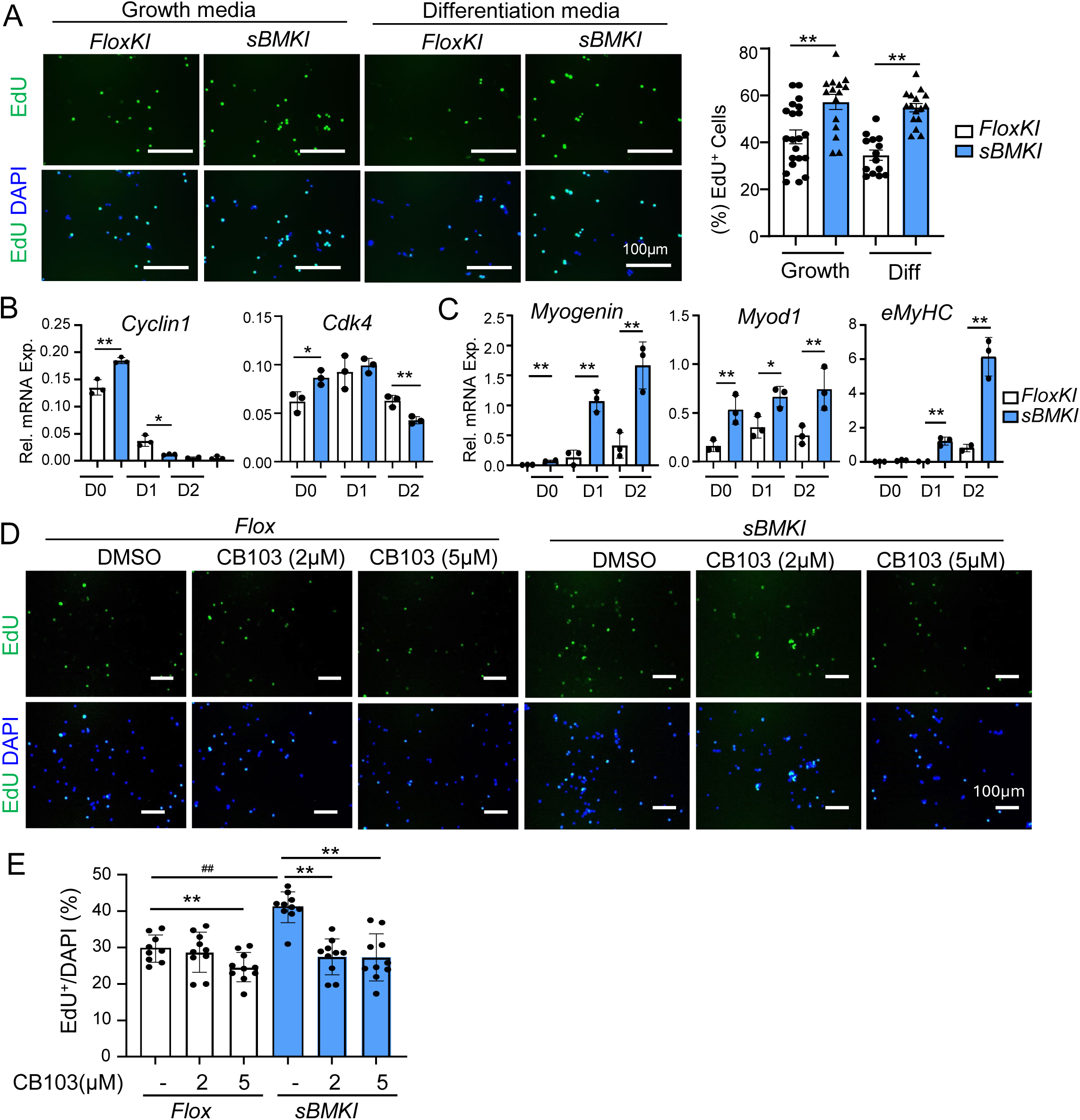
Cell-autonomous effect of Bmal1 on promoting myogenic progenitor proliferation. (A) Representative images of EdU staining with quantitative analysis of Flox and sBMKI primary myoblasts under proliferative growth condition and following one day of differentiation. (B, C) RT-qPCR analysis of dynamic expression of components of cell cycle regulators (B) and myogenic factors (C) in Flox and sBMKI primary myoblasts before and during myogenic differentiation for 2 days. N=3/group. * and **: p<0.05 or 0.01 by Student’s t test. (D, E) Representative images of EdU staining (D) with quantitative analysis (E) to determine effect of the Notch inhibitor CB103 on blocking Flox and sBMKI primary myoblasts proliferation under proliferative growth condition. *, **: p<0.05 or 0.01 CB103 vs. DMSO control, and ##: Flox control vs. sBMKI.by one-way ANOVA with Tukey’s post-hoc analysis.

### Pharmacological activation of clock by chlorhexidine and CM002 promotes myogenesis via Notch and Wnt signaling

We recently identified a clock activator, chlorhexidine (CHX), that displayed strong pro-myogenic propertiesvia activation of Wnt signaling^25^. Based on the chemical scaffold of this hit molecule, we performed medicinal chemical modifications to generate a series of analogs followed by a biochemical screen of clock-activating properties together with functional assays to determine their pro-myogenic activities. This led to the discovery of a new derivative CM002 that displayed increased clock-modulatory activity as a lead compound (**Fig. 5A & S5**). In C2C12 myoblasts, both clock-activating molecules were sufficient to induce the expression of identified Notch pathway genes along with activation of Wnt signaling as indicated by *Axin 2* level (**Fig. 5B**). CM002 displayed an overall modest increase in its efficacy on inducing these clock targets. Furthermore, CM002 treatment of myoblasts was sufficient to promote clock activity, as shown by the significantly enhanced Bmal1 occupancy of clock target promoters within the Notch pathway (**Fig. 5C**). The fold of induction stimulated by CM002 was promoter-specific, with Notch 1 highly responsive resulting in a ∼34-fold induction as compared to basal Bmal1 occupancy on this promoter. This finding promoted our investigation into the potential of CM002 in eliciting circadian time-dependent clock activation of Notch target genes, using synchronized myoblasts with CM002 treatment to determine its modulation of Notch genes. Following serum shock synchronization, a circadian oscillation of Bmal1 protein was induced, with its peak abundance occurring between CT17-23. We thus compared Notch gene induction by CM002 treatment at this circadian time window as compared with its trough period at CT29-35. As shown by the induction of Notch transcripts, including Notch 1, Noth 3 and Jag 1, CM002 treated at ZT17 resulted in markedly higher levels than the treatment at ZT29 that induced minimal response (**Fig. 5D**). In comparison, expression of *HeyL* revealed a distinct different response, with robust CM002 inductions at both time points. As a effector of Notch signaling, this response of *HeyL* could be potentially due to its function in integrating upstream Notch activity altered by CM002. Consistent with CHX and CM002 activity in inducing clock-controlled Notch signaling, both compounds were able to stimulate the proliferation of primary myoblasts to a largely similar degree (**Fig. 5E**).

**Figure 5.**
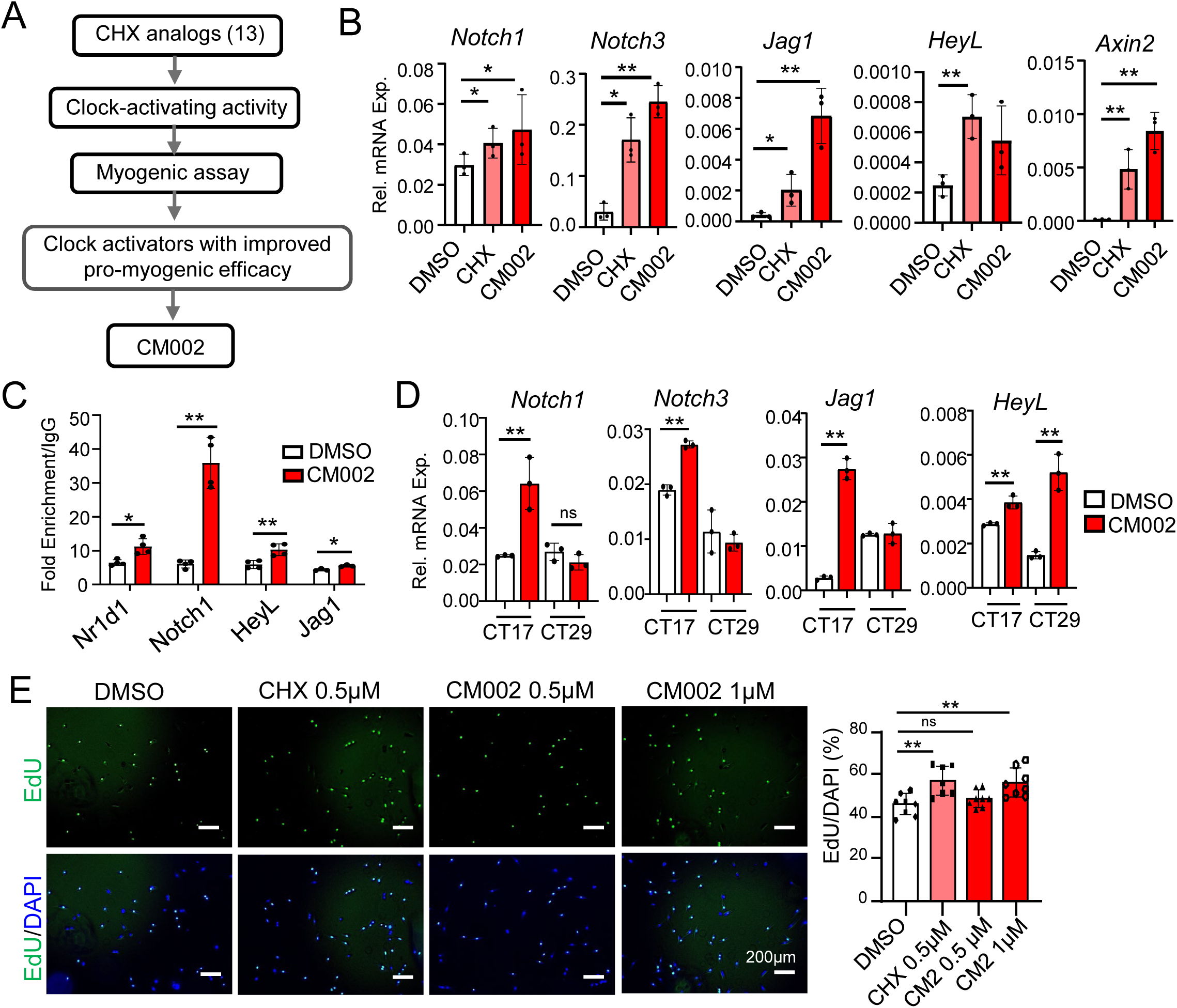
Effect of clock-activating molecules on inducing Notch and Wnt pathway. (A) Schematic flow chart of generating analogs of chlorhexidine leading to the screen and identification of CM002 as a novel clock-activating lead compound. (B) RT-qPCR analysis of Notch and Wnt pathway genes of C2C12 myoblasts treated with CHX (0.5µM) or CM002 (0.5µM) for 6 hours. * and **: p<0.05 or 0.01 CM002 or CHX vs. DMSO control by Student’s t test. (C) ChIP-qPCR analysis of chromatin occupancy of Bmal1 on identified target promoters involved in Notch signaling. C2C2 myoblasts were treated with vehicle (0.1% DMSO) or 0.5µM CM002 for 6 hours prior to chromatin extraction (n=3). * and **: p<0.05 or 0.01 CM002 vs. DMSO control. (D) Effect of time-dependent effect of CM002 on modulating Notch signaling components by RT-qPCR analysis in synchronized myoblasts. Serum shock by one hour treatment of 20% serum was used for synchronization. Cells were treated with DMSO or CM002 (0.5uM) for 6 hours prior to sample collection at CT17 and 29 after serum shock. *, **: p<0.05 or 0.01 CM002 vs. control by one-way ANOVA with Tukey’s post-hoc analysis. (E) EdU fluorescence labeling to assess CHX and CM002 effect on primary myoblast proliferation under normal growth conditions. (n=7-8/group). *, **: p<0.05 or 0.01 CM002 vs. DMSO by one-way ANOVA with Tukey’s post-hoc analysis.

### Clock activation promotes the myogenic differentiation of mouse and human dystrophic myoblasts

Given the function of clock in promoting myogeneis and its regulation of the Wnt signaling, we determined whether clock-activating molecules modulate myogenic differentiation. Immunofluorescence staining of myosin heavy chain (MyHC), a mature myocyte marker, demonstrated that 0.5 µM CHX markedly enhanced mature myofibers formation after 3 days of differentiation, as compared to DMSO control (**Fig. 6A**). MyHC^+^ myofibers induced by CM002 at 0.5 µM was stronger than that of CHX with further increase observed at 1 µM, suggesting a stronger efficacy in stimulating myogenic differentiation. When treated in primary myoblasts isolated from mdx mice, a preclinical model for DMD, CHX was sufficient to promote the differentiation of dystrophin-deficient myoblasts, as indicated by MyHC+ myofiber staining 3 days after of myogenic induction (**Fig. 6B**). Furthermore, in Bmal1-deficient mdx primary myoblasts obtained from BMKO/mdx double-null mice^33^, this effect of CHX was completely abolished, indicating the clock-dependency of CHX effect on promoting myogenesis. Similar to the higher efficacy of CM002 than that of CHX in promoting myogenesis in C2C12 myoblasts, CM002 demonstrated a stronger effect in primary myoblasts as well, as indicated by increased abundance and differentiated morphology of MyHC^+^ myotubes (**Fig. 6C**). Lastly, to explore the potential for clock-targeting approaches to augment regenerative capacity in dystrophic disease, we tested these compounds on myogenic differentiation of primary myoblasts derived from patients with Duchenne Muscular Dystrophy (DMD). In human DMD primary myoblasts, both CHX and CM002 were sufficient to induce up-regulation of Bmal1 protein together with markedly elevated levels of myogenic factors MyoD and myogenin, suggesting activation of the myogenic response with comparable levels of inductions (**Fig. 6D & 6E**). These findings collective suggested the potential of pharmacological clock activation to enhance myogenesis for DMD intervention.

**Figure 6.**
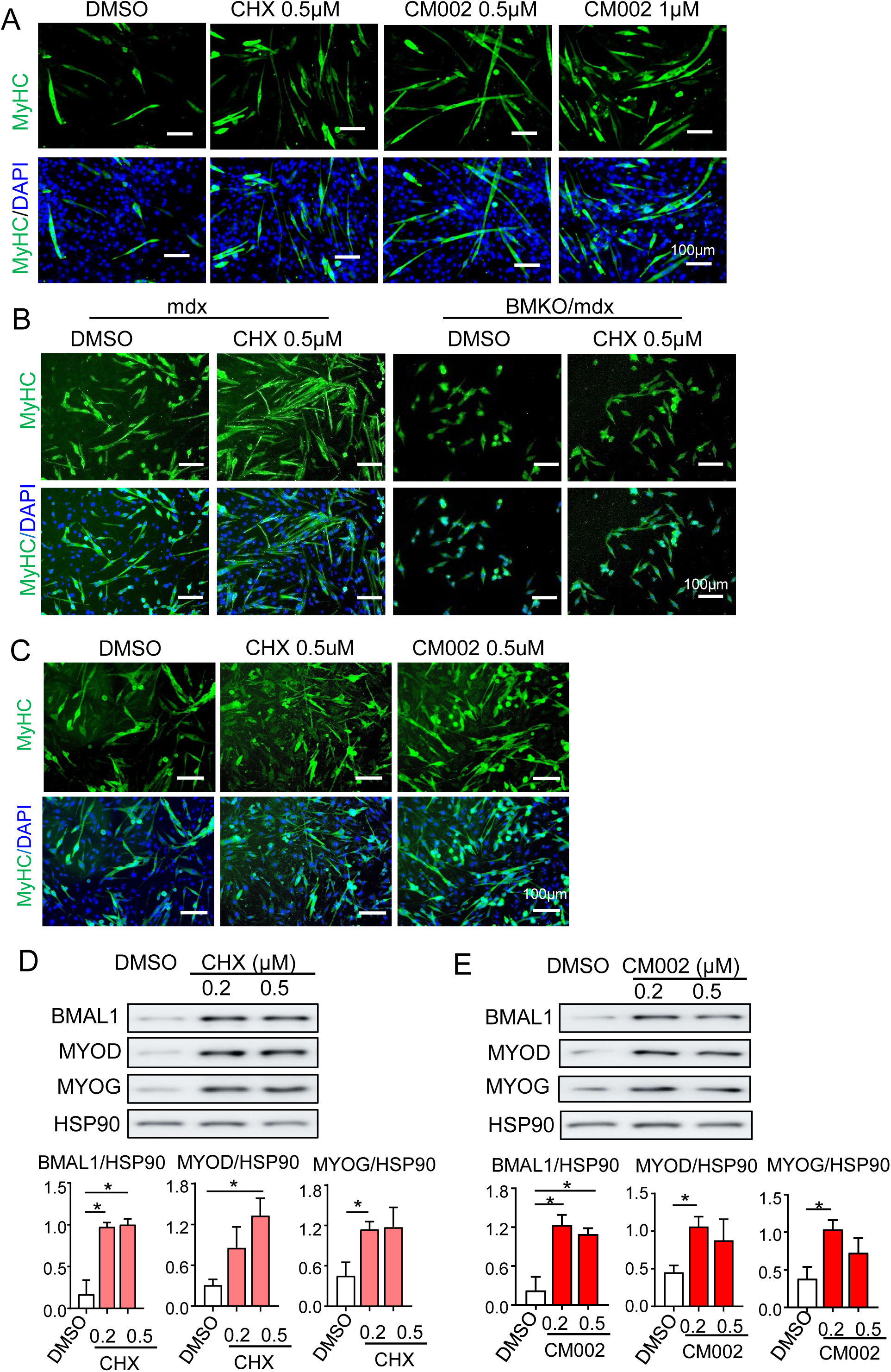
Clock activators promote myogenic differentiation of mouse and human dystrophic myoblasts. (A) Representative images of immunofluorescence staining of myosin heavy chain (MyHC, green) of C2C12 myoblast myogenic differentiation at day 4 with chlorhexidine (CHX) or CM002 treatment at indicated concentrations. (B) Representative images of MyHC staining of differentiation of primary myoblasts derived from mdx and Bmal1-deficient mdx mice (BMKO/mdx) treated with DMSO or CHX (0.5 μM). Primary myoblasts were differentiated for 3 days using 2% horse serum. (C) Representative images of MyHC staining of mdx primary myoblasts differentiation for 3 days with 0.5 μM of CHX or CM002 treatment. (D, E) Representative images of immunoblot analysis of the effect of CHX (D) and CM002 (E) on clock and myogenic gene regulation using human primary myoblasts with DMD following 3 days of differentiation. Quantitative analysis is based on three independent repeats. *p<0.05 and ** p<0.01 CM002 or CHX treatment vs. DMSO control by Student’s t test.

### Clock activators display pro-myogenic activities upon acute muscle injury and in chronic dystrophic milieu

Given the pro-myogenic effects of clock-activating molecules, we next explored their potential for enhancing regenerative repair in vivo in response to acute muscle injury or during chronic damage in the dystrophic disease milieu. Four consecutive intramuscular deliveries of CHX or CM002 into the TA muscle, administered one day after cardiotoxin injury, were used to determine their effect on regenerative myogenesis (**Fig. 7A**). Regeneration via eMyHC immunofluorescence staining was assessed at dpi 5 together laminin staining (**Fig. 7B)**. Vehicle-treated controls displayed low levels of new myofiber, as expected of this stage of neomyofiber formation upon cardiotoxin injury. In comparison, both CHX and CM002-treated muscles demonstrated substantial amount of eMyHC staining, suggesting sustained nascent myofiber regeneration. Interestingly, laminin staining revealed marked increase in the abundance of myofibers observed per section area in CHX or CM002-treated regenerating muscles, suggesting an enhanced regenerative response with nascent myofiber formation. As a result, myofiber size distribution at this stage revealed a robust increase in the abundance of newly-regenerated fibers with a shift toward smaller cross section area **(Fig. 7C**). In contrast, myofibers of medium size ranging from 2000-2500 μm2 were increased, suggesting an additional tendency toward accelerated regenerating myofiber size incretion at advance stages. Analysis of myogenic response using whole TA muscle revealed the elevated levels of Myf5, MyoD and Myogenein, that were more robustly induced by CM002 than that of CHX (**Fig. 7D & 7E**), though the induction of Pax7 protein was comparable. These findings, in aggregate, indicate that these clock-activating molecules are capable of promoting the regenerative response with sustained new myofiber formation following acute muscle damage.

**Figure 7.**
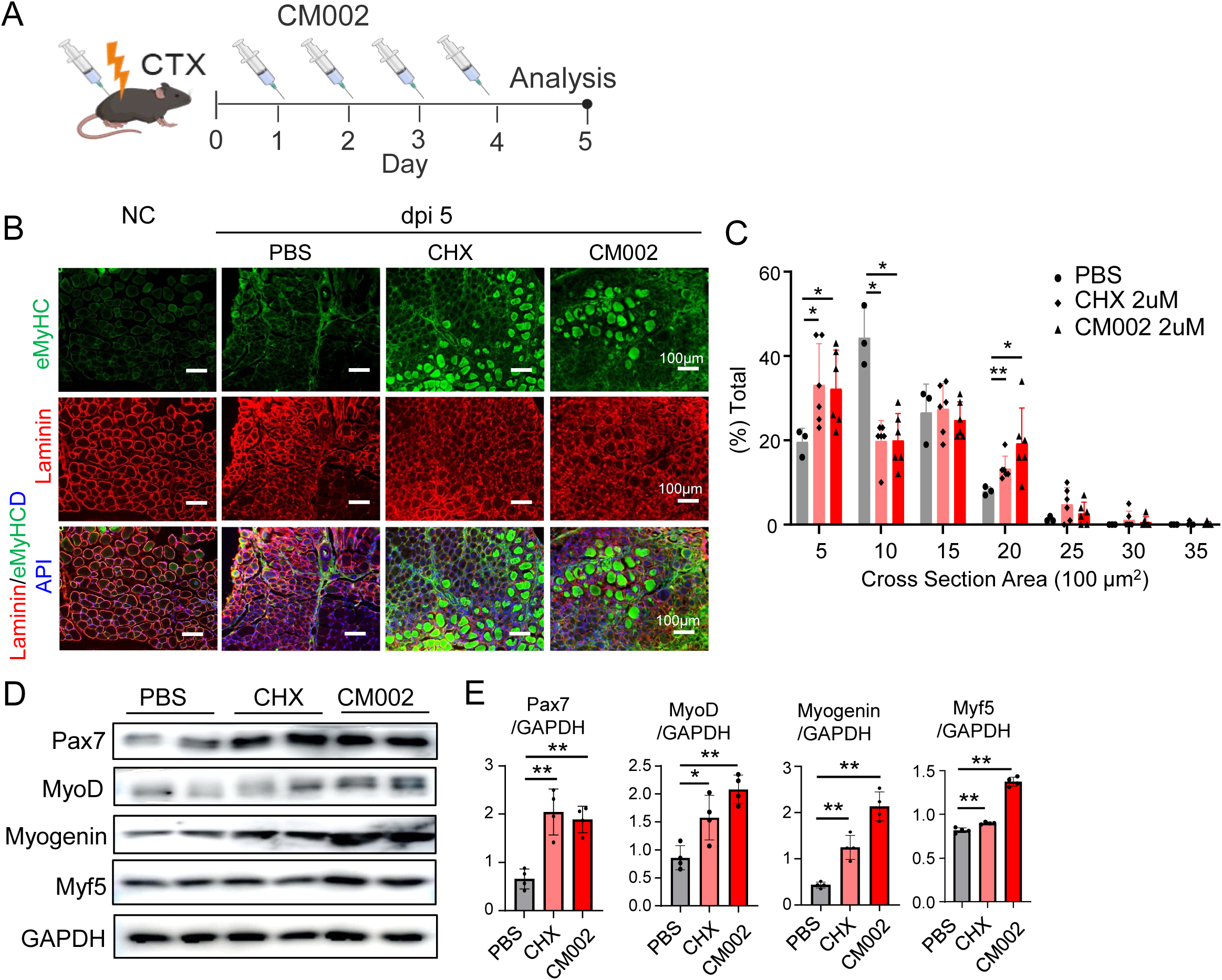
Effect of pharmacological clock activation on regenerative myogenesis upon cardiotoxin-induced injury. (A) Schematic representation of CM002 treatment time course following CTX injury of TA muscle. One day after CTX injury, 50ul of CHX (0.5uM in PBS) or CM002 solution (0.5uM in 0.1% DMSO) was injected into injured TA for 4 days with samples collected at dpi 5 for analysis. (B, C) Representative immunofluorescence staining analysis of embryonic myosin heavy chain (eMyHC) and laminin to examine new myofiber synthesis in TA muscle at dpi5, with quantitative analysis of myofiber cross section area distribution using laminin-stained images (C). (D, E) Immunoblot analysis of genes involved in regenerative myogenesis following injections with CHX or CM002 in CTX-injected TA muscle, with quantitative analysis using three repeats (E). Each lane represents pooled samples of 4-6 mice/group, with quantitative analysis from three repeats.

Next, we tested whether the effect of CHX and CM002 on promoting regeneration could be of potential applications for muscular dystrophy using the mdx mice. CHX or CM002 were administered following a similar design of CTX-induced regeneration via local delivery into the TA muscle (**Fig. 8A**). In the dystrophic muscle, these compounds elicited strong inductions of key clock regulators, Bmal1, CLOCK and its direct target gene Rev-erbα (**Fig. 8B**). Interestingly, CM002 displayed stronger effect on inducing clock activation in mdx muscle than that of CHX. Furthermore, these molecules also resulted in robust up-regulations of key signaling components of the Notch and Wnt pathways, as demonstrated by qRT-PCR analysis of identified direct targets of Bmal1 (**Fig. 8C**) indicative of activation of clock output pathways. Protein levels of cleaved Notch, a critical mediator of intracellular Notch signaling, and the Notch pathway effector, Hes1, were increased by both compounds, further validating enhanced Notch signaling activity in treated mdx muscle (**Fig. 8D**). β-catenin protein were also increased, suggesting potential induction of Wnt pathway as well by these compounds. Consistent with their differing efficacy on inducing clock regulators, CM002 effect on these Notch pathway components were stronger than that of CHX. Additional analysis of CHX and CM002-treated muscle revealed elevated Pax7 and Cyclin D1 proteins levels, along with strong inductions of myogenic factors (Myf5, MyoD & Myogenin (**Fig. 8E**), suggesting enhanced regenerative myogenesis in the dystrophic disease milieu by both compounds with comparable efficacy. The proliferative rate of satellite cells in dystrophic muscle following treatments was assessed using EdU labelling (**Fig. 8F**). Compared to vehicle controls, the proportion of EdU^+^/Pax7^+^ satellite cells in both CHX and CM002-treated cohorts were increased by ∼52%, while the total amount of Pax7^+^ satellite cells were comparable between these groups, suggesting increased population of cycling satellite cells without significant expansion of the total pool size. Staining of nascent myofibers revealed a strong increase in eMyHC+ neo-myofiber formation in CHX or CM002-treated groups (**Fig. 8G**). Similar to the findings from CTX injury, a striking increase in the abundance of small regenerating myofibers were present in these groups. This resulted in an overall size distribution shifting toward smaller myofibers and a corresponding reduction of larger sizes, indictive of the emergence of small nascent myofiber following treatments (**Fig. 8H**). Collectively, these in vivo effects of clock-activating molecules on augmenting regenerative repair in the mdx muscle suggest their potential applications in dystrophic disease that may help to restore the impaired regenerative capacity.

**Figure 8.**
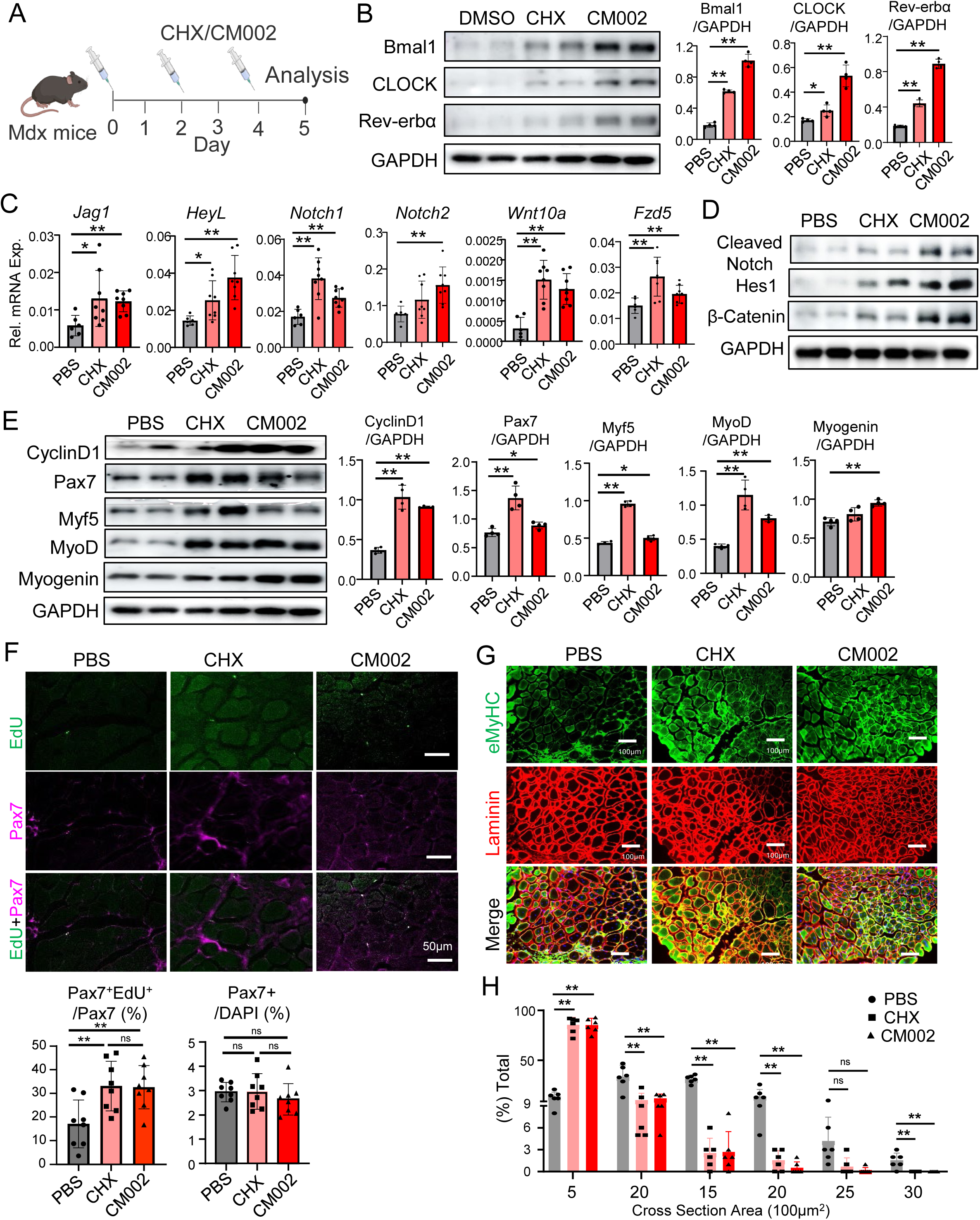
Effect of CM002 and CHX on promoting regenerative myogenesis in dystrophic muscle. (A) Schematic representation of CHX and CM002 treatment time course in mdx mice. Male mdx mice were treated every other day with these compounds (50 μl of 2μM in PBS) for a total of 3 times via intramuscular injection into TA with samples collected after 5 days. (B) Representative images of immunoblot analysis of clock regulators in TA muscle of mdx mice treated with CHX or CM002 with quantitative analysis from three repeats. (C, D) RT-qPCR (C), and immunoblot analysis (D), of Notch and Wnt signaling components in TA muscle of mdx mice with CHX or CM002 treatment. (E) Representative images of immunoblot analysis of regenerative myogenesis in TA of mdx mice treated with CHX or CM002. Each lane represents pooled samples of 3-4mice/lane. (F) Representative images of immunofluorescence Pax7 staining (Magenta) with EdU labeling (green) to determine proliferation in TA of mdx mice. N=8/group. (G, H) Representative images of immunofluorescence staining of embryonic MyHC (green) and laminin (red) of mdx mice TA cross section following CHX or CM002 treatments (G) with quantitative analysis (H). *p<0.05 and ** p<0.01 CM002 or CHX treatment vs. DMSO control by Student’s *t* test.

## Discussion

The immense regenerative potential of muscle stem cells could be exploited for muscle disease applications^34,35^. Despite mechanistic investigations of circadian clock function in orchestrating regenerative myogenesis, targeting clock modulation for muscle disease interventions has yet to be explored. By leveraging a new clock gain-of-function genetic model and pharmacological clock-enhancing interventions, our current findings demonstrated, for the first time, that enhancing muscle stem cell clock activity was sufficient to augment their regenerative properties in acute injury and dystrophic conditions. Furthermore, we identified that direct target genes of clock among the key components of Notch signaling pathway, and that the coordinated control of a Notch and Wnt activity by clock promotes proliferative expansion and myogenic progression during regenerative myogenesis.

To date, most of the studies of circadian clock function in skeletal muscle stem cells are limited to loss-of-function models. By generating a model with Bmal1 over-expression in satellite cells, our current study demonstrated the pro-myogenic potential of augmenting clock activity specifically via modulation of the properties of myogenic precursor cell population. Notably, clock gain-of-function in MuSCs does not lead to overt phenotypes in skeletal muscle under basal conditions. In addition, a surprising finding was that upon acute injury, the peak of regeneration in *sBMKI* mice was initially delayed 3 days after injury, though a later peak at dpi5 was augmented with an elevated and prolonged regenerative myogenic time course. This regenerative dynamic correlated with an early induction of the Notch signaling with enhanced Wnt pathway activity in later time points of regeneration. While current findings revealed that the Notch pathway is also under clock transcriptional control with Bmal1 chromatin occupancy detected at E-box sites of identified gene promoters, we have reported previously that key signaling components of Wnt cascade were direct clock targets^5,30^. During muscle regeneration, a coordinated temporal transition from activation of Notch at the early stage to Wnt signaling drives myogenic progression^19,29,34,36^. By activating a clock-controlled intrinsic temporal regulatory control conferred, either genetic or pharmacological activation of the clock facilitates a normal regenerative timing-appropriate transition from Notch to Wnt pathway that enables proliferative expansion of the MuSC pool with coordinated myogenic progression. The Notch pathway is known to play a crucial role in driving the self-renewal to maintain the satellite cell pool^29,37^ with its activation displaying context- and developmental stage-dependent effects on myogenesis^38,39^. The increased Notch activation in sBMKI MuSCs in early stages of regeneration may have extended proliferative expansion phase with maintenance of self-renewal, and inhibition of Notch signaling in these cells indeed blocked the increase in their proliferation. Given the oscillatory nature of molecular clock action via a negative feed-back loop, Bmal1 promotion of Notch signaling and consequent MuSC modulations are likely not persistent, and thus may account for the distinct effect as compared with constitutive Notch activation in satellite cells^37,38^. It is also

Notch signaling is dominant during the early phase after muscle injury or during muscle stem cell maintenance. The Notch signaling, a key pathway in mediating neighboring cell-cell interactions, is known to confer cell-autonomous cis-inhibitory effect on signaling within the same cell, while activating Notch intracellular signal cascade of the receiving cell via trans-activation^40–42^. In myogenesis, a recently identified mechanism of MyoD activation of the Notch ligand, Dll1, is involved in inhibiting Notch signaling via cis-inhibition^43^. Based on findings of the target genes of clock involved in Notch signaling pathway, there are potential mechanisms that may account for the self-limited activation of this pathway in sBMKI mice during regeneration. Jag-1, a known Notch ligand, was found to be a direct clock target. Regulation of this ligand in sBMKI cells, known to induce a cis-inhibition mechanism in the originating cell itself while stimulating neighboring receiving cells via transactivation, may elicit distinct signaling outcomes that poses inhibition of this pathway during regeneration. A cis-inhibition mechanism mediated by Jag1-Notch 1 in sBMKI-expressing committed myogenic progenitors may lead to the attenuation of initial Notch with myogenic progression, while allowing nearby MuSC to maintain a self-renewal pool with activation. It is also possible that, as HeyL was also a target of clock regulation, this repressive transcriptional effector of Notch as well as mediating a negative feedback loop, may contribute to the prolonged but eventual dampening of Notch activation in and thus the myogenic drive at the end of regeneration induced by clock activity in sBMKI mice. Thus, by re-enforcing an intrinsic temporal regulatory element involved in distinct signaling cascade required for normal regenerative myogenesis, MuSC clock activation augments the overall myogenic response that facilitates the orderly progression through distinct phases of myogenic activation, stem cell expansion, myogenic commitment and differentiation. Besides Notch and Wnt regulations, circadian clock was reported to modulate various developmental signaling pathways to control the homeostatic functions of various stem cell compartments, including the Tgf-β and BMP pathways^2,44^. Thus, it remains to be investigated whether additional clock-controlled signaling mechanisms could be involved in its fine-tuning of MuSC behaviors that may constitute a potential intricate network of temporally controlled signaling events involved in muscle regeneration, or potentially other tissue remodeling processes.

Our current study is the first report in elucidating clock function in controlling both Notch and Wnt activities to facilitate normal myogenesis. During regenerative myogenesis in sBMKI mice, potential interactions between Notch and Wnt signaling driven by clock gain-of-function may have contributed to a prolonged proliferative phase of satellite cells in early stages with a delayed transition nascent myofiber formation. Notch signaling was known to be involved in satellite cell activation and proliferation^22,29^. To further direct that enhanced Notch activity early during regeneration mediated the increased proliferative expansion of satellite cells, we used primary myoblasts obtained from sBMKI satellite cells and found they indeed display a higher proliferative rate that was efficiently blocked by inhibition of Notch signaling. Thus, clock-controlled Notch activation underlies the proliferative phenotype in sBMKI cells, similarly to our prior findings of impaired Wnt signaling in mediating the defect in differentiation of Bmal1-null myoblasts^5^. In addition, clock activator CM002 enhanced chromatin occupancy of Bmal1 on E-boxes of Notch gene promoter regions, providing direct support of enhanced Notch activity by pharmacological clock activation. In comparison, satellite cell-specific overexpression of the Notch Intracellular Domain, the mediator of Notch signaling following its protease cleavage, resulted in a block of myogenic development leading to loss of muscle mass^37^. Activation of Notch is known to block myogenic differentiation by inhibiting the expression of myogenic factors^45–47^. The intriguing finding of a delayed but prolonged regenerative time course in the sBMKI could be due to the over-activations of Notch at early stages, while subsequent activation of Wnt pathway was sufficient to facilitate the myogenic transition with consequent positive impact on regeneration as demonstrated by a moderate shift toward larger myofiber size. Thus, likely due to the dual regulation of clock on Notch as well as Wnt, and potentially other pathways, the impact of clock gain-of-function on satellite cells is distinct from that of promoting Notch activity.

Discovery of clock-activating molecules enabled our study of pharmacological clock interventions in muscle regeneration under normal and dystrophic disease conditions. Chlorhexidine (CHX) as a novel circadian clock activator with pro-myogenic activities^25^, and we explored its potential applications in muscular dystrophy using primary myoblasts derived from the mdx mice as a cellular model with further demonstration of its clock-dependent actions. Consistent with findings of Notch and Wnt pathway are direct transcriptional targets of clock, Bmal1 overexpression or clock activation by CHX or CM002 were able to induce both Notch and Wnt pathway genes in myogenic precursors. Importantly, clock regulation of Notch and Wnt pathways were applicable to the dystrophic disease condition, with CHX and CM002 treatment in mdx mice resulting in induction of these signaling components. Thus, clock-controlled Notch and Wnt signaling activation may mediate, to a significant extent, the efficacy of the pharmacological activators of clock on promoting regenerative myogenesis in vivo, suggesting these potentials of clock regulation could be pharmacologically targeted to promote intrinsic mechanisms that orchestrate regenerative myogenesis. Both Wnt and Notch signaling abnormalities were found in the dystrophic muscle that contributes to impaired regenerative capacity that is a key pathogenic mechanism of loss of muscle mass and function^48–50^. Current findings of CHX and CM002 on enhancing regeneration using the acute injury model provided the proof-of-principle evidence that clock activation could be leveraged to promote regenerative repair. More importantly, in the dystrophic mdx model, we demonstrate that clock activation induced both the Notch and Wnt pathways, leading to increased satellite cell proliferation and new myofiber regeneration in this chronic disease condition. These findings of clock gain-of-function interventions on orchestrating MuSC behavior provide the basis for targeting clock function to promote regenerative capacity for disease interventions, and the ability of CHX and CM002 in augmenting regenerative response under dystrophic condition further their therapeutic development. The chromatin occupancy of CLOCK/Bmal1 displays circadian oscillation^51,52^ and clock activators, as we demonstrated, can promote clock chromatin occupancy on target promoters. Therefore, a potential time-dependent in vivo efficacy of pharmacological activation of clock warrants further investigations toward chronotherapeutic applications.

Collectively, our current proof-of-concept studies demonstrated the potential to develop clock-targeting interventions for muscle disease treatments. Nonetheless, significant hurdles remain to ultimately progress toward clinical drug development and application. This study is limited to intramuscular delivery with the goal to test initial in vivo effects. Due to the current scope, whether these clock-activating molecules could be applied for systemic delivery to promote regenerative capacity in mdx mice with functional impact on the disease development trajectory remains to be seen. Detailed pharmacokinetic studies and extensive toxicity testing are needed to enable systemic delivery effort. This study thus remains as the first step toward future systematic in vivo investigations. Based on the chemical scaffold of these clock-activating molecules we’ve developed, additional high-affinity drug-like leads and ultimately clinically efficacious drugs with desirable safety profiles can be synthesized, tested and for therapeutic applications. Therefore, our current study established the mechanistic basis to target the clock function to augment regenerative myogenesis and provide the first proof-of principle testing to boost regenerative capacity to ultimately help patients with DMD.

---

**Figure S1.**
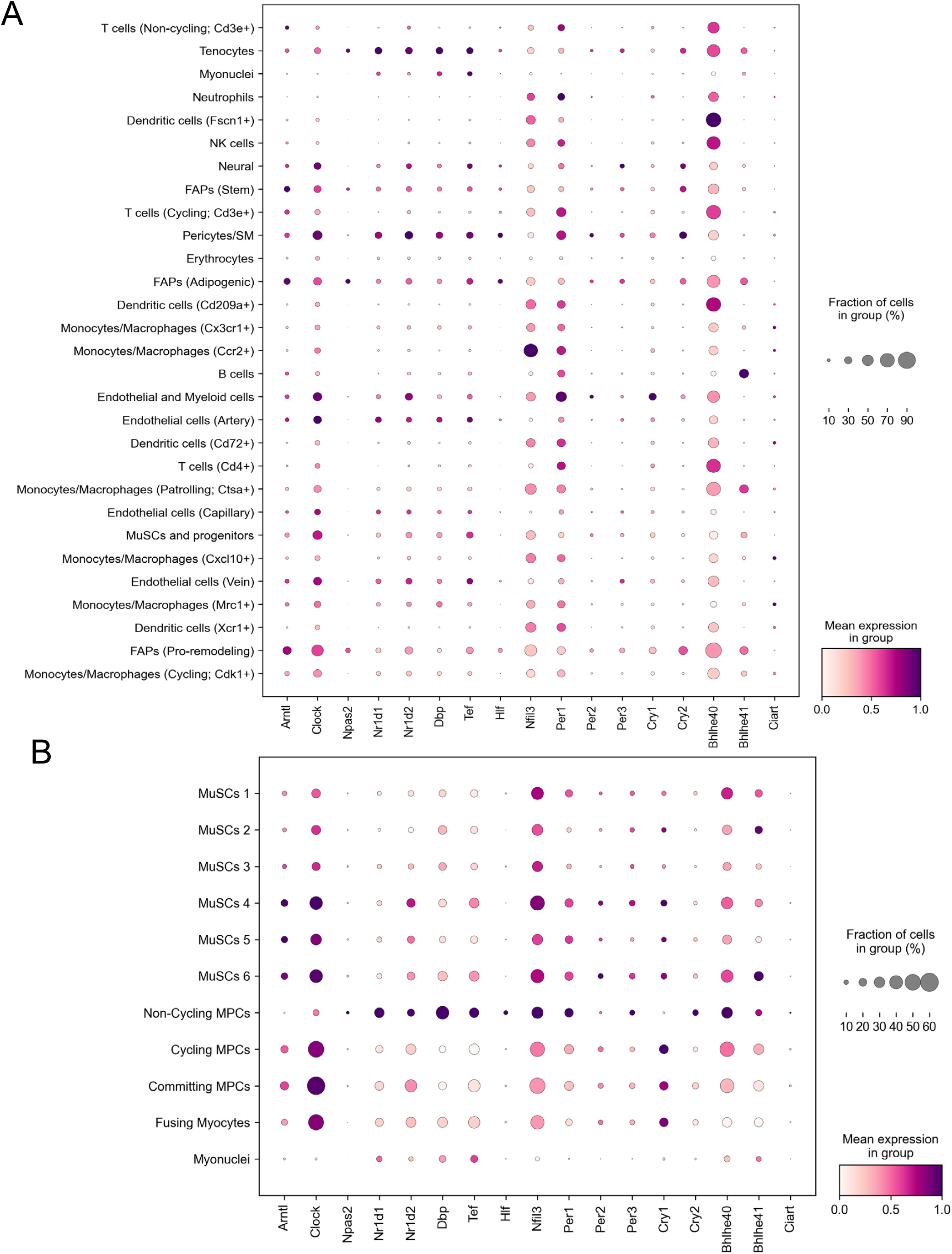
Clock gene expression in distinct cell types identified in sc-RNA seq analysis of muscle regeneration. (A) Clock gene expression in all cell types involved in regeneration using previously published sc-RNA-seq datasets of muscle regeneration with dpi 0 to 7. (B) Clock gene expression in subgroups of MuSCs and myogenic progenitor cells (MPCs).

**Figure S2.**
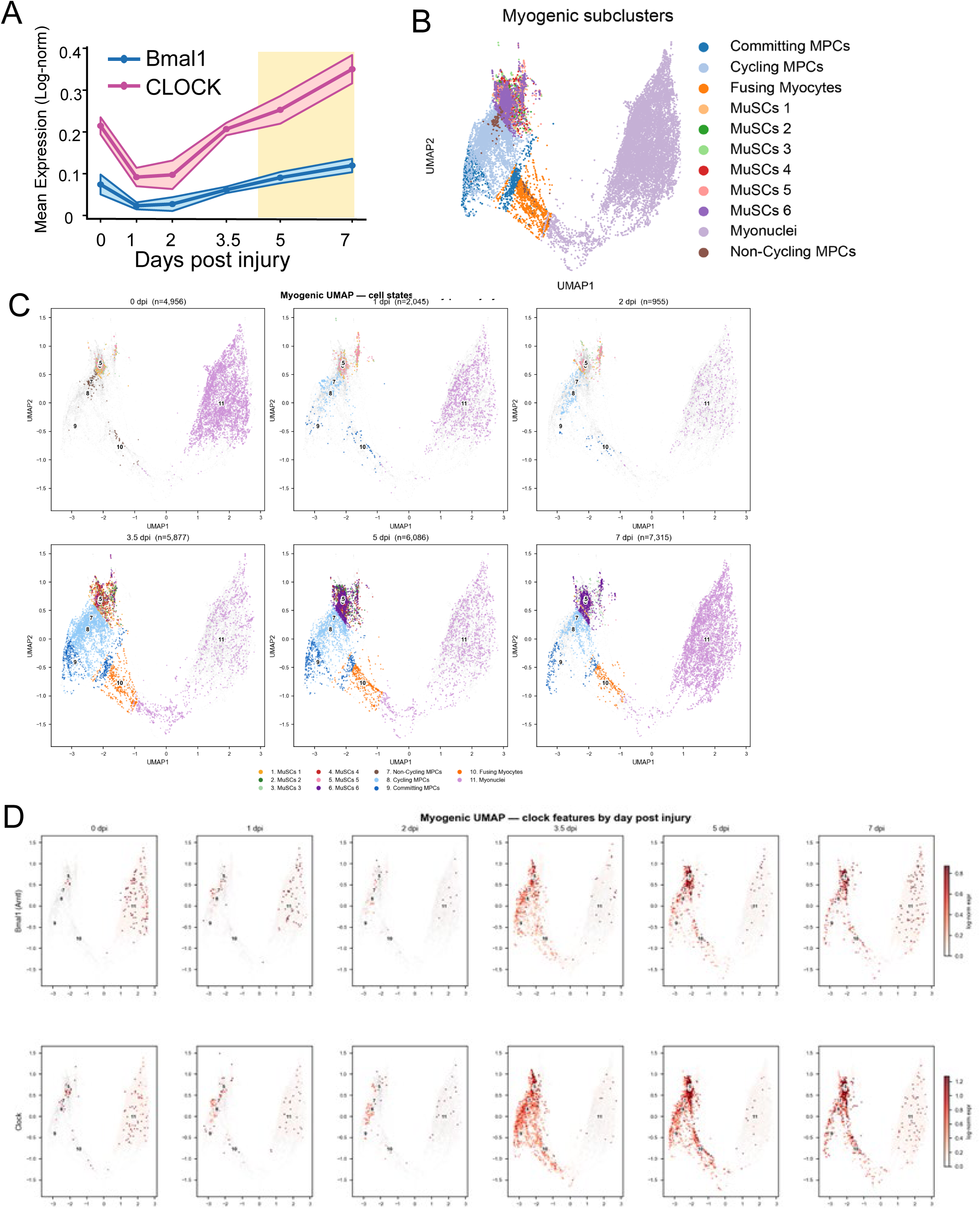
Clock gene expression dynamics across regeneration and association with Notch signaling. (A) Bmal1 and CLOCK aggregate expression in MuSCs along regeneration time course of dpi 0 to 7. (B, C) Identification of distinct myogenic subclusters (B) and the changes across dpi 0 to 7. (D) Bmal1 and CLOCK in subclustered of MuSCs and MPCs along regeneration.

**Figure S3.**
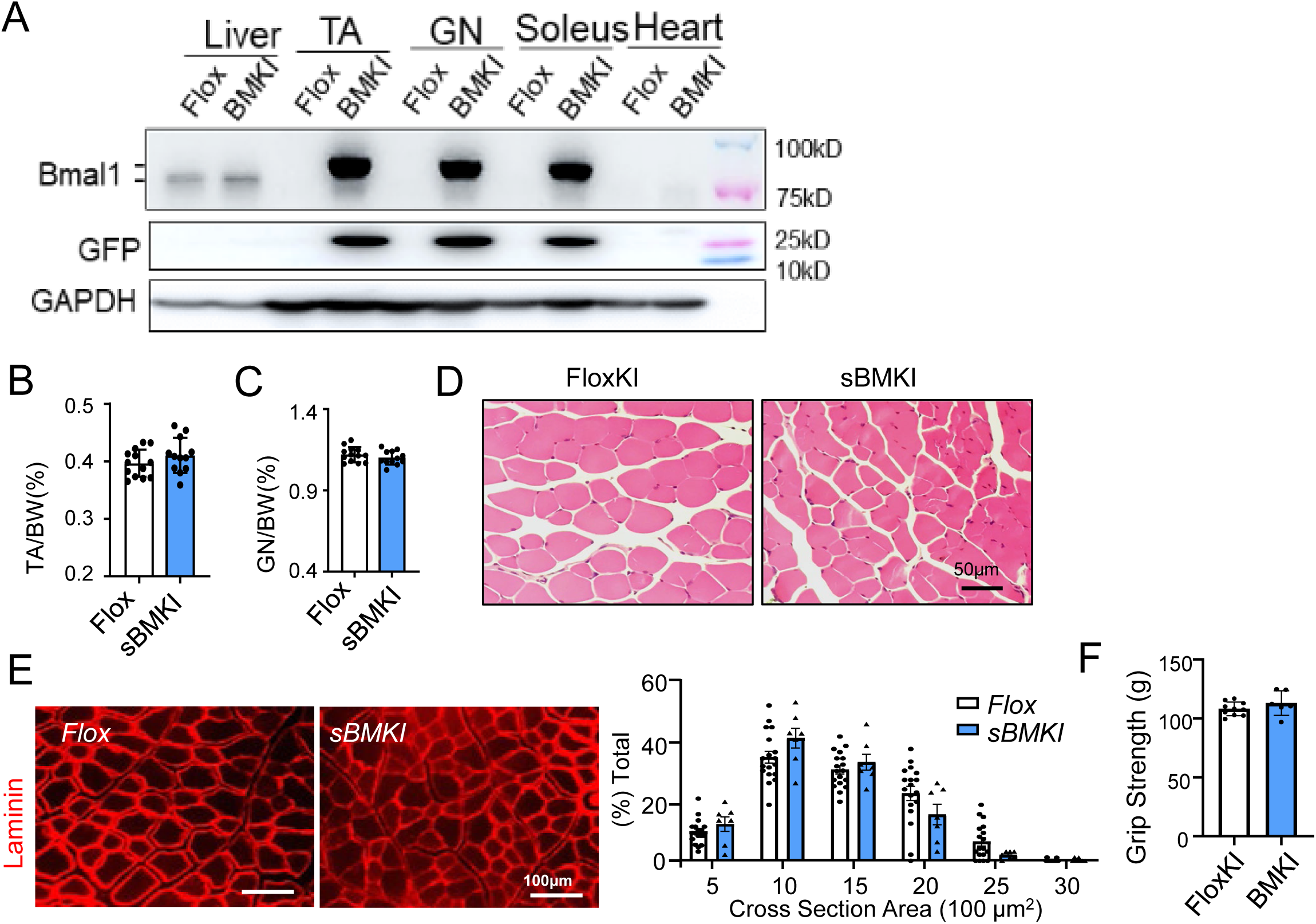
Basal characterization of satellite cell-specific Bmal1 overexpression in mice. (A, B) Validation of satellite cell-specific Bmal1 overexpression model. (A) Representative immunoblot analysis of Bmal1 (A) in distinct muscle in mice with satellite cell-specific overexpression (sBMKI) and floxed knock-in controls (Flox). Exogenous Bmal1 knockin protein ∼95 kD as shown in sBMKI muscles. TA: Tibialis Anterior; GN: Gastrocnemius. Each lane represents pooled samples of n=5-6 mice/group. (B, C) Analysis of tibialis anterior (TA, B) and gastrocnemius muscle (GN, C) weight normalized by body weight in 3-month-old male Flox control and sBMKI mice. N=10-12/group. (D) Representative H/E histology of TA cross section in Flox control and sBMKI mice. (E) Representative images of immunofluorescence staining of laminin to assess of cross section area of TA muscle of Flox control and sBMKI mice, with quantitative analysis of size distribution. (F) Analysis of hanging forelimb grip strength test of male Flox and sBMKI mice (N=6-7/group).

**Figure S4.**
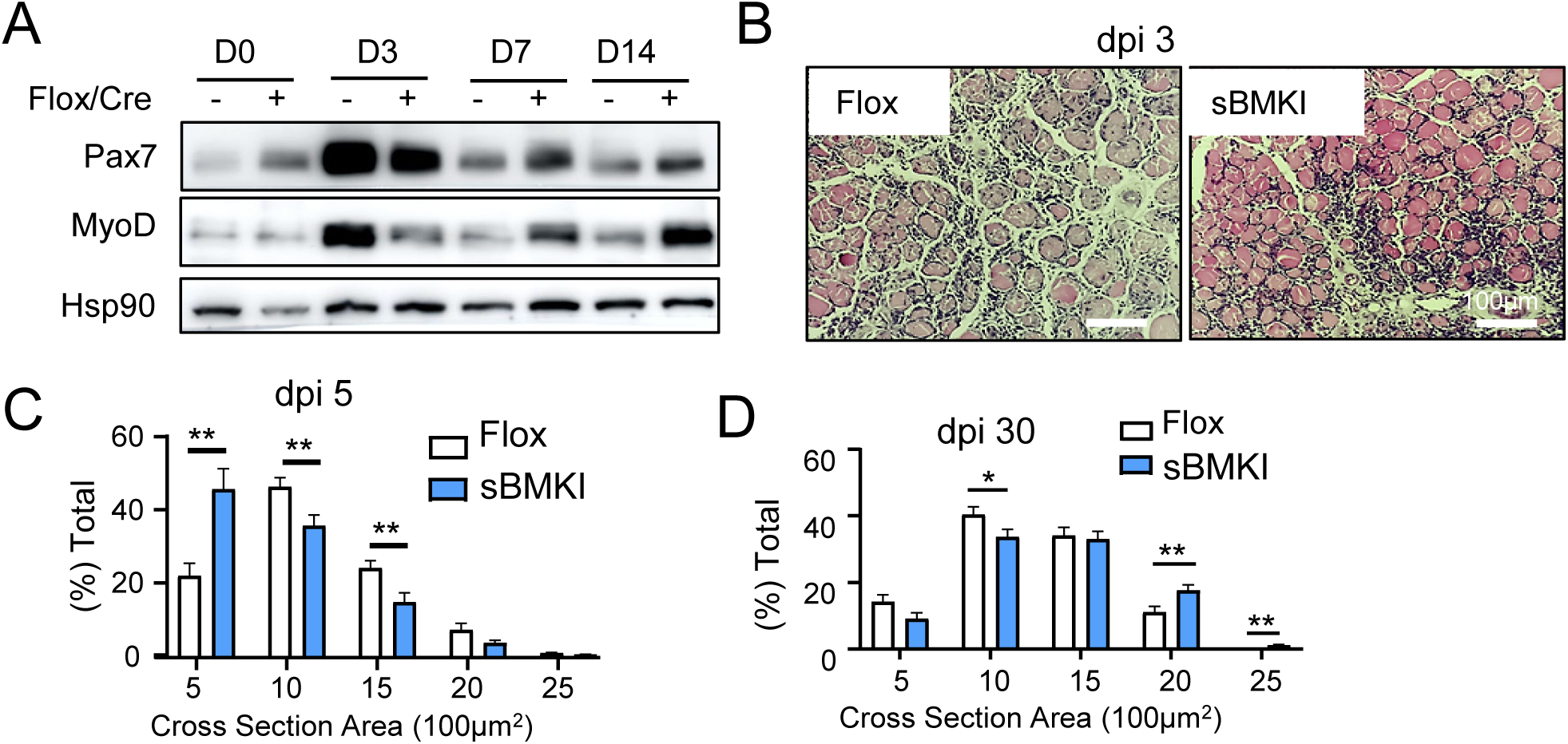
Analysis of muscle regeneration in mice with satellite cell-specific Bmal1 overexpression. (A) Representative images of immunoblot analysis of Pax7 and MyoD expression level at indicated time points till 14 days following cardiotoxin-induced injury in TA muscles of male Flox and sBMKI mice. N=6-8 mice/group. Each lane represent a pool of 6-8 mice. (B) Representative images of H/E histology of Flox and sBMKI at day 3 after CTX injury. (C, D) Analysis of myofiber cross section area distribution at dpi 5 and dpi 30 in Flox and sBMKI mice. N=6-8 mice/group.

**Figure S5.**
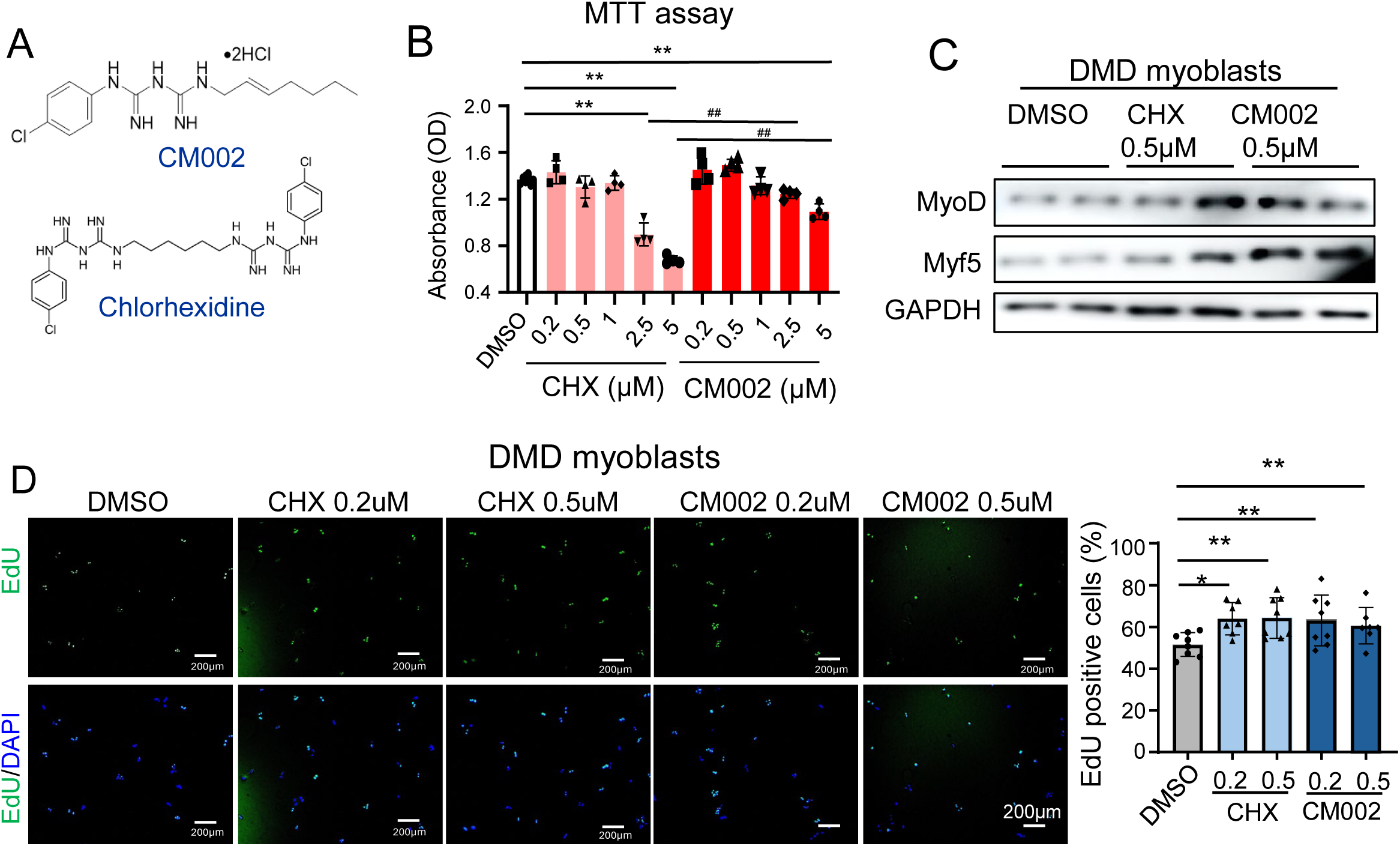
Effect of clock-activating molecules on viability and myogenic properties of myoblasts. (A) Chemical structure of clock activating molecules Chlorhexidine (CHX) and CM002. (B) MTT assay to assess viability of primary myoblasts treated with CHX or CM002 at indicated concentrations. N=3/group. (C) Representative images of immunoblot analysis of myogenic factors of human myoblasts with DMD treated with CHX or CM002. Human DMD myoblasts were differentiated for 3 days (D) Representative images of EdU incorporation to assess proliferation of C2C12 myoblasts treated at indicated concentrations of CHX or CM002 for 4 hour with quantitative analysis. N=7-8/treatment group. Scale bar: 200μm.

